# Inherent envelope fluctuations in forward masking: Effects of age and hearing loss

**DOI:** 10.1101/2022.10.20.513056

**Authors:** Marc A. Brennan, Adam Svec, Afagh Farhadi, Braden Maxwell, Laurel H. Carney

**Affiliations:** University of Nebraska-Lincoln, Lincoln, NE 68583; San José State University, San Jose, CA 95192; University of Rochester, Rochester, NY 14642

**Author notes:** Correspondence: Marc Brennan University of Nebraska-Lincoln 4075 East Campus Loop Lincoln, NE 68583.

**Keywords:** aging, hearing loss, forward masking, auditory model

## Abstract

Forward masking is generally greater for Gaussian noise (GN) than for low-fluctuation noise maskers, i.e., GN disruption. Because the minimal hearing loss that is associated with older age may affect GN disruption differently than more significant hearing loss, the current study explored the contribution of minimal hearing loss associated with older age to GN disruption. GN disruption was measured using three masker-signal delays (25, 75, and 150 ms) for three adult groups: younger participants with normal hearing, older participants with minimal hearing loss, and older participants with sensorineural hearing loss. The role of underlying mechanisms was tested using a computational model for midbrain neurons. The primary result suggests that older listeners with mild threshold elevations that typically occur with age may be more susceptible to the deleterious effects of masker-envelope fluctuations than younger listeners with normal hearing. Results from the computational model indicate that there may be a larger influence of efferent feedback and saturation of inner hair cells on forward masking and GN disruption than previously thought.

## I INTRODUCTION

The temporal envelope of Gaussian noise (GN) contains short-term fluctuations in level. As the bandwidth of GN decreases, these fluctuations tend to increase the amount of masking for both simultaneous maskers (Pumplin, 1985; Zurek and Durlach, 1987; Hartmann and Pumplin, 1988; Kohlrausch *et al*., 1997; Savel and Bacon, 2003) and forward maskers (Moore, 1981; Svec *et al*., 2015, 2016) relative to a masker of equivalent bandwidth and level, for which the temporal envelope fluctuations have been minimized, referred to here as low-fluctuation noise (LFN). Given that less forward masking is generally expected and observed for LFN than for GN (e.g., Svec *et al*., 2015, 2016), the additional masking attributed to the envelope fluctuations is often assessed by subtracting the LFN threshold from the GN threshold, referred to here as GN disruption. Considering that masker fluctuations negatively affect speech recognition (Stone *et al*., 2011, 2012), the mechanisms that contribute to GN disruption have implications for understanding the processes that contribute to the speech recognition difficulties often experienced by listeners in noisy situations (Füllgrabe *et al*., 2015; Brennan *et al*., 2016). The purpose of this study was to assess plausible mechanisms that contribute to GN disruption as a function of age and hearing status. To accomplish this, GN disruption at three different masker-signal delays was assessed for younger normal hearing (YNH), older participants with minimal hearing loss (OMHL), and older participants with sensorineural hearing loss (OSNHL).

### A Cochlear Compression and Listener Uncertainty

Cochlear compression and listener uncertainty could play a role in GN disruption. For participants with sensorineural hearing loss (SNHL), a linearized cochlear response should yield larger effective amplitude peaks than a compressive cochlear response (i.e., as for YNH); the linearized cochlear would thus likely yield greater GN disruption. These larger effective amplitude peaks may result in an increased potential for confusing an amplitude fluctuation near the end of the masker for the onset of the brief signal, referred to as listener uncertainty or the “confusion effect” (Moore, 1981; Moore *et al*., 1985; Neff, 1986). However, such an explanation alone cannot 1) account for the greater GN disruption for a 25 ms masker-signal delay that occurred for OMHL relative to both YNH and OSNHL participants by Svec *et al*. (2015) and 2) the greater GN disruption observed by Svec *et al*. (2016) for OSNHL relative to YNH participants for a 75-ms masker-signal delay—a delay time for which confusion effects should no longer occur. As elaborated below, one potential reason for the greater GN disruption for the OMHL participants in Svec *et al*. (2015) was that these participants had slightly higher hearing thresholds than the YNH participants.

### B Inhibition at the Inferior Colliculus

Multiple peripheral and central processes, such as the middle-ear muscle reflex, neural adaptation, efferent innervation from the medial olivocochlear bundle, and dynamic range adaptation could also contribute to forward masking and, by extension, to GN disruption (See a review by Jennings, 2021). Nelson *et al*. (2009) argued that single auditory-nerve fibers (ANFs) tuned to the frequency of the signal cannot account for forward masking due to a limited dynamic range of discharge rates and maximum threshold shifts, two attributes of ANF responses (in anesthetized animals) that do not mirror psychophysical forward-masking results. After measuring recovery from forward masking at the level of the inferior colliculus (IC) in awake marmosets, Nelson and colleagues asserted that the phenomenon of forward masking is likely a consequence of inhibitory neural responses arising between the auditory nerve and the IC. Based on thresholds estimated for both GN and LFN, recent physiological work has suggested that GN disruption for tone-in-noise stimuli is observable from extracellular recordings of IC cells in awake rabbits (Fan *et al*., 2021).

### C Medial Olivocochlear Efferent System

The Medial Olivocochlear (MOC) efferent system could play a role in GN disruption. The MOC system provides efferent input to the outer hair cells (OHCs) that regulates OHC electromotility. Increased MOC activity is associated with decreased OHC electromotility and, in turn, reduced cochlear gain and smaller vibrations on the basilar membrane (Fuchs and Lauer, 2019). Upward shifts of 2-14 dB of auditory-nerve rate-level functions with electrical stimulation of the MOC have been observed (Gifford and Guinan, 1983), with the largest shifts occurring for levels between 45- and 75-dB SPL (Guinan, 2018). The MOC efferent system receives ascending inputs from peripheral auditory neurons as well as descending projections from the IC and other more central auditory structures (Mulders and Robertson, 2000; Schofield, 2011). While the cumulative contribution of the MOC to auditory physiology and perception remains unclear (e.g., Jennings, 2021), Carney (2018) argued that one purpose of the MOC efferent pathway might be to regulate OHC gain to control the saturation of inner hair cells (IHCs), thereby preserving cross-characteristic frequency (CF) contrast in low-frequency temporal fluctuations in ANF responses, referred to as neural fluctuations. Due to the fact that IC neurons are sensitive to envelope fluctuations (reviewed in Joris *et al*., 2004) and the IC has descending projections to the MOC, Carney additionally argued that neural fluctuations may excite MOC efferents and decrease OHC electromotility, leading to reduced cochlear gain. Farhadi *et al*. (2021) modeled IC firing rate as a function of time in response to amplitude-modulated stimuli. The results suggested that predictions were not accurate without the inclusion of efferent-regulated cochlear gain driven by IC inputs to the MOC system, providing additional support for the assertion that fluctuating inputs to the IC are likely directly affecting MOC spike rates, and in turn, cochlear gain. According to the computational model of Farhadi *et al*. (2021), the effect of this decreased cochlear gain in response to a fluctuating input, such as GN, should be decreased IHC saturation, and thus greater ANF fluctuations.

Decreased cochlear gain following stimulus presentation begins to decay within 25 ms of the masker offset (Roverud and Strickland, 2010), but appears to remain, in diminished form, up to 50 s following stimulus presentation (Brown, 2001; Cooper and Guinan, 2003). This initial time course of decreased cochlear gain following stimulus presentation roughly corresponds with the time course of recovery for forward masking, with an exponential decay in masking that extends from 0 to approximately 128 ms following masker offset (e.g., Jesteadt *et al*., 1982). If cochlear gain is influenced by temporal-envelope fluctuations through IC projections to the MOC, masker envelope fluctuations associated with a GN forward masker should induce a reduction in cochlear gain and a concomitant decrease in the response of the IC to a signal that follows a masker stimulus. Consequently, the decrease in masking associated with longer masker-signal delays would be expected to follow the time course of decreased cochlear gain following stimulus presentation, primarily from 25 to 125 ms (Brown, 2001; Cooper and Guinan, 2003; Roverud and Strickland, 2010; Rabbitt and Brownell, 2011). In contrast, when the IC receives relatively small fluctuations (e.g., produced by LFN), the IC projections to the MOC should induce less cochlear-gain reduction relative to a stimulus with larger fluctuations (e.g., GN). The net effects would likely result in a higher IC rate in response to a signal following a LFN masker than following a GN forward masker, which is consistent with GN disruption. These effects of the MOC efferent system could account for the gradual decrease in GN disruption observed with increasing masker-signal delay (25, 50, and 75 ms) for participants with NH (Svec *et al*., 2016).

Additional important factors affecting the MOC influence on GN disruption are: a) the *relative* degree of IHC saturation in response to the GN and LFN forward maskers; and b) changes in MOC physiology associated with age and SNHL. The relative IHC gain and saturation in response to these maskers, and thus the neural fluctuations and the MOC responses that they elicit, would be expected to vary across groups of participants. For younger participants with audiometric thresholds within normal limits, IHCs will tend to saturate in response to both GN and LFN forward maskers when presented at high sound levels, resulting in a relatively small neural fluctuation rates in response to both maskers, and thus relatively little predicted GN disruption for moderate and high-level maskers. For older participants with slightly elevated audiometric thresholds (e.g., small amounts of hearing loss), 1) the threshold of the IHC response is elevated, 2) higher displacements of the basilar membrane are needed to achieve the same IHC responses, and 3) the displacement associated with saturation is also shifted to a higher input level. Consequently, less IHC saturation would be expected and fluctuations in the ANF response to GN and LNN would differ just as the stimulus envelopes differ. Although the influence of older age on the MOC efferent system is unclear (Fuchs & Lauer, 2019), neural fluctuations and changes in cochlear gain would still be expected, which together may result in relatively large GN disruption for older participants with small amounts of hearing loss relative to participants with better hearing.

For participants with significant SNHL, two factors may affect the magnitude of GN disruption. First, even when masker presentation levels are high, IHCs might not saturate for either GN or LFN maskers due to threshold elevation and a shift towards higher input levels of the IHC response to basilar membrane displacement. Second, the impact of the MOC efferent system on cochlear gain is reduced (Carney, 2018; Fuchs and Lauer, 2019). Consequently, less GN disruption would be expected for older participants with substantial SNHL than for older participants with minimal hearing loss. After an initial cochlear-gain decrease associated with an input stimulus, an intact MOC system allows cochlear gain to increase during the interval from 25 to 125 ms or more after stimulus offset (Brown, 2001; Roverud and Strickland, 2010), implying that differences in GN disruption for YNH, OMHL, and OSNHL should be maximal at a short masker-signal delay and decrease for longer masker-signal delays.

### D Research Questions

To clarify possible physiological mechanisms associated with GN disruption, the current study examined the effects of age and SNHL on forward-masked thresholds obtained using maskers with relatively large (GN) or relatively small (LFN) fluctuations in level over time. Masked thresholds were obtained for a 4000-Hz pure-tone signal when presented after the offset of GN or LFN maskers at three masker-signal delays (25, 75, and 150 ms) for YNH, OMHL, and OSNHL participants. Contributions of inhibition and the MOC efferent system to GN disruption were evaluated using a computational model of midbrain neurons. The following hypotheses were formed:

1) If a lack of cochlear compression strongly contributes to GN disruption (e.g., by increasing the effects of uncertainty), then GN disruption should be greatest for the OSNHL participants, less for OMHL participants, and least for YNH participants.
2) If differences in the physiological response of the MOC efferent system for GN and LFN maskers strongly contribute to GN disruption, then GN disruption should vary non-monotonically with audiometric threshold at the signal frequency. The most GN disruption is hypothesized to occur for participants with relatively small amounts of hearing loss (OMHL participants), for whom IHC saturation and neural fluctuations would differ the most between the two masker types. Additionally, a computational model incorporating the MOC efferent system should better predict GN disruption than a model without the MOC efferent system.

For all mechanisms that contribute to GN disruption, differences in GN disruption for each group were hypothesized to decrease as the masker-signal delay increased.

## II METHODS

### A Participants

A total of 57 participants were enrolled in this study. Five participants were excluded because they did not qualify based on their hearing thresholds (see below). Of the remainder, 18 YNH (19 – 25 years, mean (M)=22), 14 OMHL (65 – 81 years, M=70), and 20 OSNHL (62 – 82 years, M=71) participants completed the study. Within each group, the age of the participants was evenly distributed across their respective age ranges. A graduate student in audiology measured hearing thresholds (ANSI, 2004) for all participants using conventional audiometry (ASHA, 2005) at 6000 Hz and at octave frequencies from 250 to 8000 Hz. Hearing thresholds for the test ear are plotted in Fig. 1. Hearing thresholds for the non-test ear were within 15 dB of those for the test ear as measured by pure tone average (500, 1000, 2000 Hz). NH was defined as hearing thresholds less than 30 dB HL from 250 to 4000 Hz. However, one OMHL participant with a hearing threshold of 30 dB HL at 500 Hz participated. SNHL was defined as air conduction thresholds from 35 to 65 dB HL at 4000 Hz and, for frequencies with hearing loss greater than 25 dB HL, bone conduction thresholds within 10 dB of the air conduction threshold. Participants who did not have NH, SNHL outside the range of 35 to 65 dB HL at 4000 Hz, or a 25 dB or greater difference in air-conduction thresholds between the two ears at 4000 Hz did not qualify. All subjects were native English speakers. Data were collected at the University of Nebraska-Lincoln. Approval for this study was obtained from the Institutional Review Board. Participants consented to join the study and were paid for their time.

**Fig. 1.**
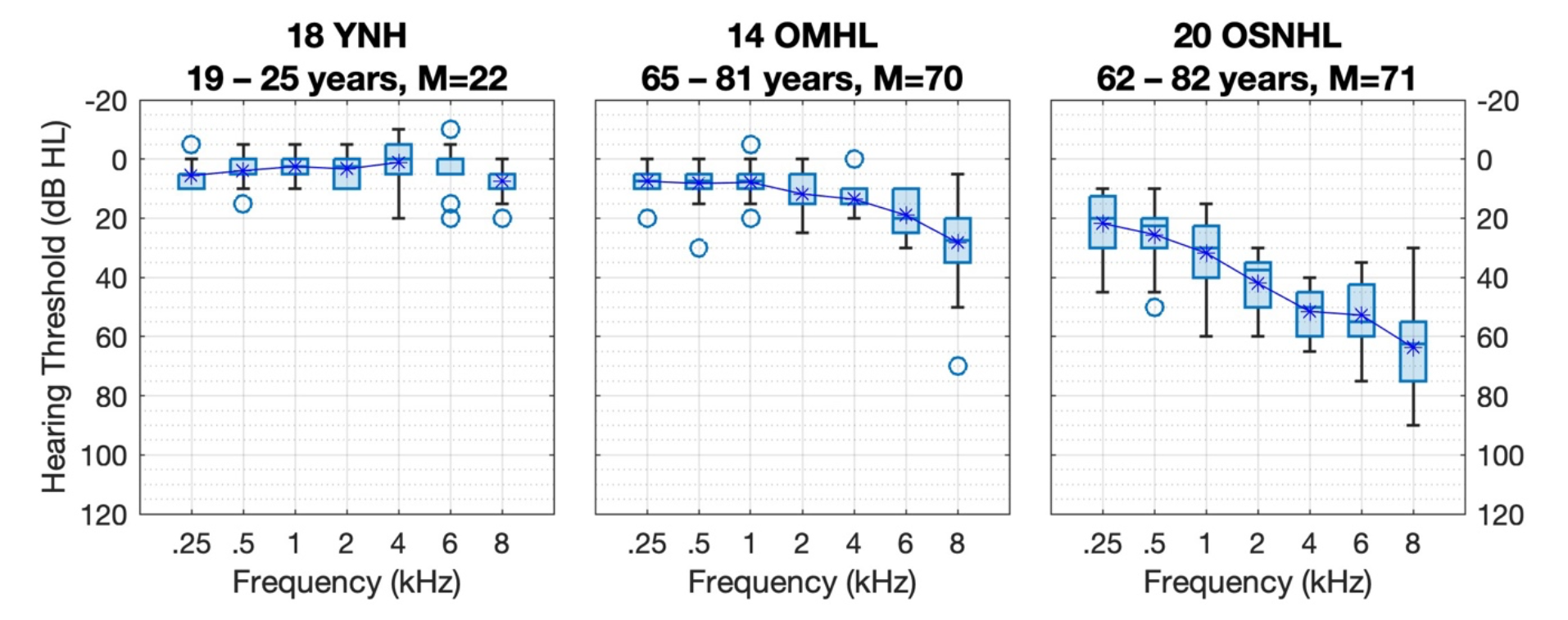
(color online) Plots of hearing threshold for each participant group. Number of participants, age range, and mean (M) age are also provided. For this and the remaining box and whisker plot, each box represents the interquartile range, each line represents the median, each asterisk represents the mean, each circle represents an outlier (>1.5 times the interquartile range), and whiskers represent the most extreme value that is not an outlier.

### B Stimuli and Apparatus

Stimuli were generated using a personal computer (22.05 kHz sampling rate; 24 bits per sample) and custom MATLAB (2019a) scripts. All durations were measured from the 0- amplitude points of the onset and offset ramps. Using the formula provided by Glasberg and Moore (1990), the masker bandwidth was one-third equivalent rectangular bandwidth (1/3 ERB_N_) centered at 4000 Hz (Glasberg and Moore, 1990). Maskers were 400 ms in duration, including 5-ms cosine-squared ramps. LFN was generated using a Hilbert transform (Buss *et al*., 2006) with the PsyAcoustX package (Bidelman *et al*., 2015). Specifically, the Hilbert envelope was computed for each 1/3 ERB_N_ GN generated. Each GN was divided in the time domain by its Hilbert envelope and then multiplied by the original spectrum in the frequency domain; these last two steps were repeated ten times to yield the final version of LFN.

Stimuli were calibrated using a 6-cc flat plate coupler and a Larson Davis System 824 sound level meter (Depew, New York). Signal levels are reported as the root-mean-square peak equivalent (0.707 x peak pressure) in dB SPL. The signal was a 4000-Hz pure tone, 10 ms in duration including 5-ms cosine-squared ramps. The signal was presented either without the masker (quiet condition) or with the signal onset 25, 75, or 150 ms after the masker offset (masker conditions). Each stimulus interval was preceded by 400 ms of silence and then followed by 400 ms of silence and, for each condition, interval durations were equivalent. The 24-bit digital stimuli were converted to analog (RME Babyface sound card, Haimhausen, Germany), amplified by a HeadAmp 4 Pro headphone distribution amplifier (Baton Rouge, Louisiana), and presented to one ear using Sennheiser HD-25 headphones (Wedemark, Germany). All testing took place in a single-walled sound attenuated room, with each participant sitting in front of a touch-screen monitor.

### C Procedures

For all experimental conditions, the signal level was varied adaptively to estimate the threshold corresponding to 71% correct with a two-down, one-up rule (Levitt, 1971). For all masker conditions, the masker level was fixed at 80 dB SPL. To ensure audibility of the signal for the OSNHL participants, the starting level of the signal was 80 dB SPL. The initial step size of 18 dB was reduced to 9 dB after the first reversal and 6 dB after the second reversal. Then, a step size of 3 dB was used. Data collection ended after a total of nine reversals. The minimum signal presentation level was −10 dB SPL, and the maximum signal presentation level was 90 dB SPL. None of the participants had tracks with 3 presentation levels in a row at 90 dB SPL (i.e., none were at ceiling). The average of the levels at the last four reversals was taken as the threshold.

A trial consisted of three observation intervals separated by 300 ms. Each interval was marked by a separate button that was illuminated on the touchscreen monitor for the stimulus duration including the encapsulating silence of 400 ms, for approximately 1200 ms per interval. For the absolute-threshold condition, one randomly selected interval contained the signal. For the masker conditions, one randomly selected interval contained the masker and signal, and the other two contained only the masker. Participants indicated the interval which they believed contained the signal by pressing the corresponding button and feedback was provided. For each condition, threshold was measured three times and the final threshold was recorded as the average across the three measurements. Threshold for the absolute-threshold condition was measured first followed by the masker conditions in random order.

### D Computational Models

Computational models for auditory-nerve (AN) and midbrain responses were used to provide insight into mechanisms that might contribute to GN disruption. The computational models are publicly available for download at https://DOI.org/10.17605/OSF.IO/AJVWD. All models included stages for the cochlea, auditory-nerve fibers, and inferior colliculus (Zilany et al., 2014; Nelson and Carney, 2004). One model was an updated version of Zilany and colleagues’ model that also included MOC efferents (Farhadi *et al*., 2021). Figures 2 and 3 provide overviews of the Farhadi *et al*. model, where firing rates are depicted for one trial of each masker type for a hearing threshold of 15 dB HL at 4 kHz (corresponding to a typical OMHL participant).

**Fig. 2.**
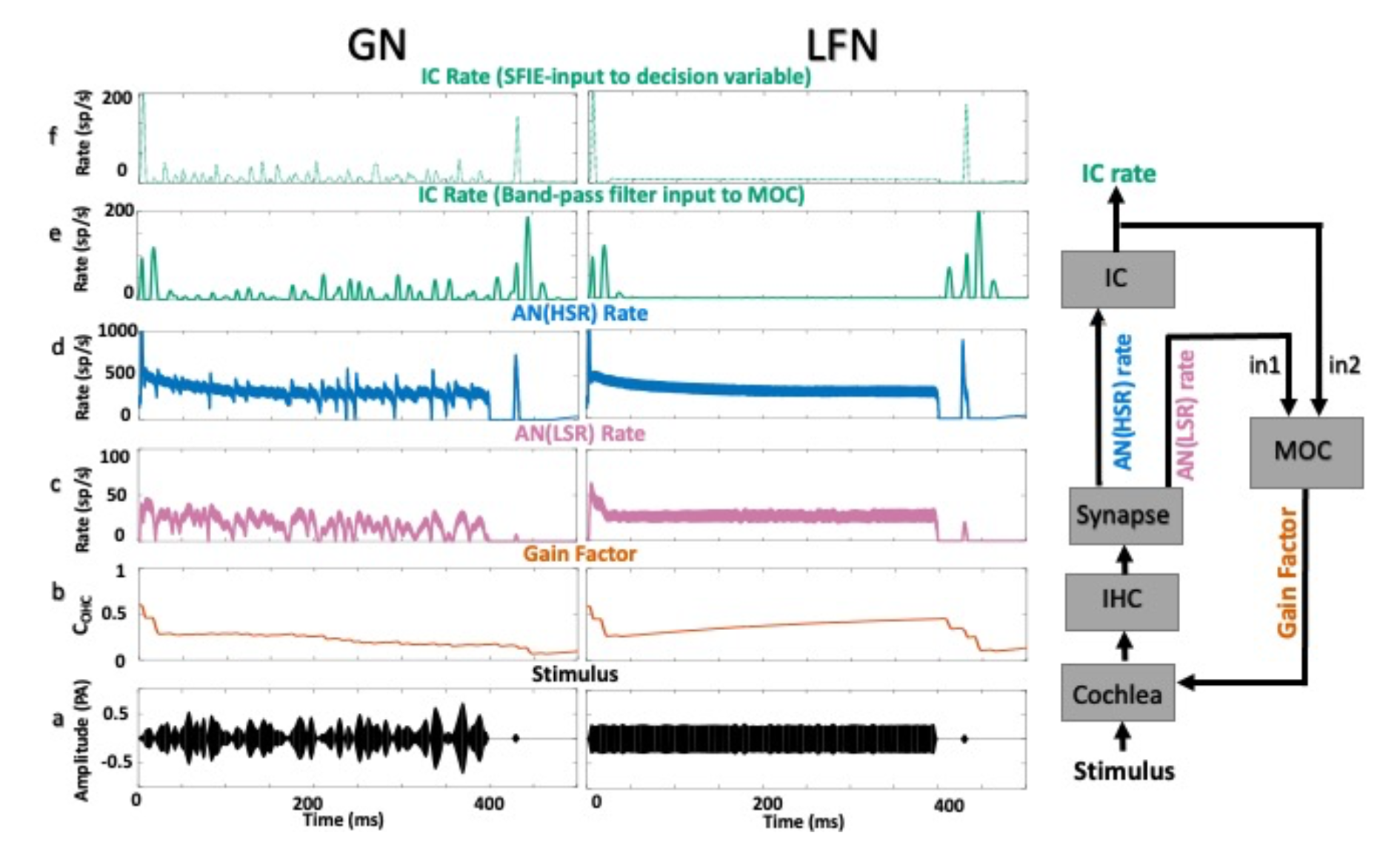
(color online). Computational model including MOC efferent system, showing example simulation of one frequency channel for OMHL. a) GN and LFN stimuli at the input to the model ultimately result in b) cochlear OHC gain factors that vary throughout the stimulus. c) Low-spontaneous rate (LSR) and d) high-spontaneous rate (HSR) AN responses are the instantaneous-rate functions at the output of the model synapse (right). e) Model IC response of band-pass filter model. The LSR fiber was used to represent wide-dynamic-range responses from the cochlear nucleus, which were combined with the IC model response as inputs to the MOC stage (right). Larger fluctuations in the IC response to GN stimuli resulted in greater gain reduction throughout the GN masker as compared to the LFN masker. Therefore, the peak rate in response to the signal for the HSR fiber was lower after the GN masker than after the LFN masker. e) Subsequently, for the IC same-frequency inhibition-excitation (SFIE) response that was the input to the decision variable, the relative rates of the signal and masker were smaller for GN than for to LFN. CF = 4 kHz. The signal level was 70 dB SPL.

**Fig. 3.**
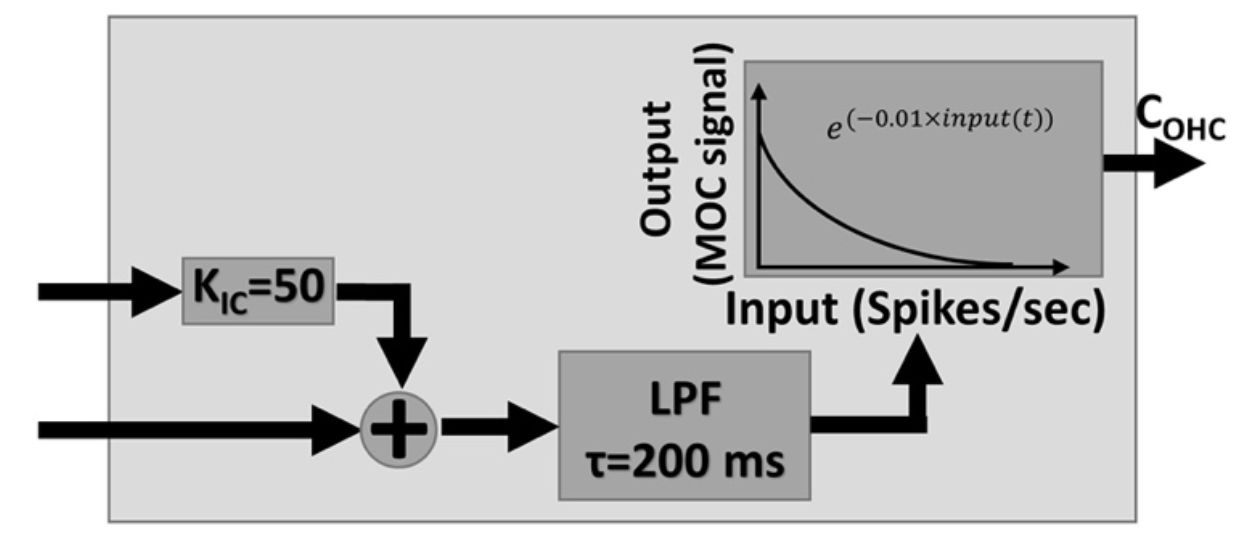
Computational model of MOC efferent system, depicting mapping of input spike rate to MOC gain factor to cochlea. Input spike rate from IC was scaled and added to that from a LSR auditory-nerve fiber, low-pass filtered, and then converted to MOC gain factor. Higher MOC spike rate resulted in smaller gain factor.

Both sets of computational models (with and without efferents) incorporated physiological properties including the saturation of IHC transduction and synaptic saturation and adaptation. Peripheral tuning of these models was based on data for human participants (Ibrahim and Bruce, 2010). Both computational models included lowpass filters corresponding to the outer and inner hair cells, each with adjustable scaling factors. Decreasing the value of the computational model parameter C_OHC_ (max=1, min=0) reduced cochlear gain, while decreasing the value of C_IHC_ reduced sensitivity of the IHC and pushed the saturation point of the IHC nonlinearity to higher input levels. The C_OHC_ and C_IHC_ scaling factors were set to match the mean audiometric threshold in dB HL for each participant group. Two-thirds of the threshold shift was ascribed to OHC, based on acoustic trauma observed in cats (Bruce *et al*., 2003; Zilany and Bruce, 2007) and estimated OHC integrity in humans with SNHL (Plack *et al*., 2004).

The model that included the MOC efferents (Farhadi *et al*., 2021) incorporated implementations of low-and high-spontaneous rate fibers (LSR, HSR) in different roles. Because the input to the ascending pathway to the IC is dominated by HSR fibers (Carney 2018), the HSR fibers provided input to the brainstem/IC model, which in turn provided one of two descending neural inputs to the MOC. The other input to the MOC consisted of afferent LSR fibers, reflecting the physiological observation that inputs to the MOC have wide dynamic ranges (Ye *et al*., 2000). Together, these two inputs to the MOC can increase cross-CF contrast in fluctuations (Farhadi *et al*., 2021). The two inputs to the MOC stage of the model were scaled to have approximately equal firing rates (the IC input was multiplied by 50), as otherwise the afferent input of the LSR fibers would dominate and limit modulation of OHC gain to enhance neural fluctuations (Figure 2). The MOC stage converted the input rates to a factor that modulated the OHC gain. This conversion was based on an exponential relationship of firing rate to gain: the cochlear gain factor decreased exponentially as the combined input to the MOC stage increased. The final step in the MOC model stage was a low-pass filter with a time constant of 200 ms, chosen to approximate the overall time course of the MOC system.

For computational efficiency, a simple, bandpass modulation filter model (Mao et al., 2013) was used for the IC response that projected to the MOC in the efferent model (IC in Fig. 2e). Because the model of the IC that provided input to the MOC had a best modulation frequency of 64 Hz, regulation of cochlear gain was driven by both overall stimulus level (via the AN LSR input) and envelope fluctuations (via the IC input). This best modulation frequency of 64 Hz was selected as it was the mean of the distribution of best modulation frequencies observed by Kim et al. (2020) for rabbits. Then, for modeling listener thresholds, the final HSR AN response for both computational models was used as the input to a SFIE implementation of the IC (Nelson and Carney, 2004) with a best modulation frequency of 64 Hz. The Discussion elaborates on the potential implications of this modeling decision. The SFIE model was used for the decision variable to avoid interference between “ringing” of the simple bandpass modulation filter at the end of the masker and the subsequent response to the signal (see Fig. 2e).

As illustrated in Fig. 2b, cochlear gain decreased after masker onset (due to high firing rates at stimulus onset for both the ANFs and IC). For the GN masker, this initial reduction in OHC gain was maintained throughout the duration of the masker and, due to the sluggishness of the MOC system, for a period after the offset of the masker. For the LFN masker, this initial reduction in OHC gain reduced over time (due to a reduction in firing rate from the IC). For the IC, these effects resulted in a greater response to the signal, relative to the masker rate, in the LFN condition. In the GN condition, the IC response during the masker was greater than in the LFN condition, and the signal response was decreased relative to the LFN condition due to decreased cochlear gain (Fig. 2b). Both higher masker response and lower signal response in the GN condition contributed to GN disruption in the model thresholds, as described further below.

One issue was the selection of CFs to include in estimating model thresholds. Presumably, participants used CFs tuned near the signal frequency of 4 kHz. Due to cochlear filtering of masker-frequency components, ANFs tuned to a masker-edge frequency will have slower fluctuations (for both masker types) than ANFs tuned to the masker center frequency. For the masker-edge CFs, there will be less IHC saturation and responsiveness to the signal stimulus. Here it was assumed that participants utilized activity in fibers with CFs that extended slightly beyond the bandwidth of the masker (3.9-4.1 kHz). Specifically, AN CFs of 3.6, 3.7, 3.8, 4, 4.2, 4.3, and 4.5 kHz were simulated. The IC stage of the model inherited these CFs, and responses were summed across CFs after the IC stage before simulating listener thresholds.

Model IC responses were obtained for a 30-dB range of signal levels near an initial, rough approximation of model threshold. Signal levels were spaced 3 dB apart, and 50 trials were completed per signal level. Three independent simulations were used for the three-interval task, one of which was selected to include the signal tone. The silence preceding each interval was set to 1 second to allow the computational model to settle before stimulus onset. Under the assumption that—for the short 25-ms masker-signal delay—the participants listened from near the end of the masker to some time past the expected temporal position of the signal, the model-IC instantaneous firing rate was recorded from 375 ms after masker onset until at least 50 ms after the end of the probe tone (the decision-variable window). The start of the decision-variable window was delayed until 425 and 500 ms for the two longer masker-signal delays, respectively, and so the decision-variable window overlapped with the response to the masker only for the 25-ms masker-signal delay condition. The inclusion of the masker response in the decision variable for the shortest masker-signal delay conditions allowed the effect of listener uncertainty to be taken into account, and this measure of listener uncertainty contributed to elevating model thresholds to better approximate behavioral thresholds.

The interval with the highest maximum instantaneous rate during this timeframe was selected as the signal interval for that trial. The proportion correct was computed for each signal level, and a logistic function was then fit to each performance-intensity curve. Threshold was estimated as the level for which the curve intersected the 70.7% correct point. This process was repeated for each masker type, masker-signal delay, and participant group.

### E Analysis

All statistical models were computed using IBM SPSS version 27 or MATLAB (2021b) and the statistics and machine learning toolbox. Mean (M) thresholds, standard deviation (SD), and percentiles were calculated. Quiet conditions across participant groups (YNH, OMHL, OSNHL) were analyzed using a repeated measures analysis of variance. Post-hoc comparisons were completed using pairwise comparisons with false discovery rate adjustments (Glickman *et al*., 2014). To assess differences in masked threshold by masker type (GN, LFN), masker-signal delay (25, 75, 150 ms), and hearing status (NH, SNHL), a linear-mixed effects model with random intercepts for each participant was conducted. Effect sizes (in dB), t-test values, and probability values are reported. Reference conditions were set to YNH, 25 ms masker-signal delay, and LFN. Using YNH as the reference group allowed the determination of effects of age and degree of hearing loss by comparing threshold for YNH and OMHL (effect of minimal hearing loss and age) and OSNHL (effect of mild-to-moderate hearing loss and age). Using the 25-ms masker-signal delay and LFN as the reference conditions allowed determination of whether GN disruption decreased from 25 to 75 ms or from 25 to 150 ms.

## III RESULTS

### A Absolute Thresholds

Figure 4 depicts detection thresholds in dB SPL. As expected, absolute thresholds (150 ms panel) were lowest for the YNH participants, with progressively higher thresholds for the OMHL and OSNHL participants (*df* = 2,50, *F* = 194.2, *p*<.001). Post-hoc testing revealed that absolute thresholds were significantly higher for OMHL (*p*<.001, M=31.1, SD=6.2) and OSNHL (*p*<.001, M=65.2, SD=9.0) participants than for YNH participants (M=21.3, SD=5.4). Absolute thresholds were also significantly higher for OSNHL than for OMHL participants (*p*<.001).

**Fig. 4.**
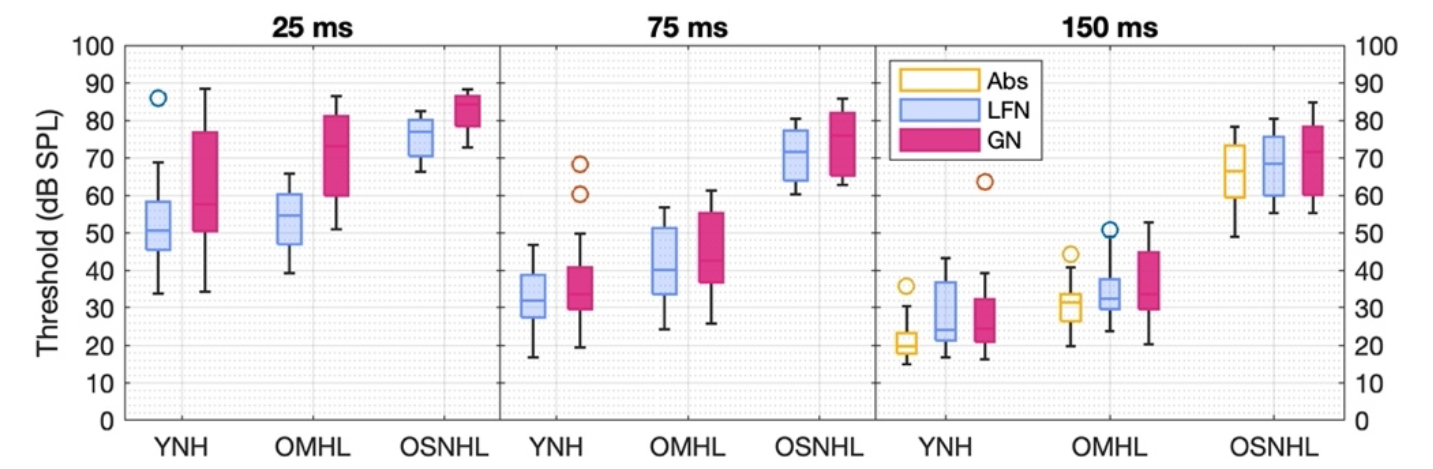
(color online) Plots of threshold with the LFN and GN maskers for each delay time. Absolute (Abs) thresholds are in the 150 ms panel. Threshold decreased as delay time increased.

### B Masked Thresholds and GN Disruption

Consistent with prior work (Brennan *et al*., 2015; Svec *et al*., 2015, 2016) and as shown in Fig. 4, masked thresholds for all conditions were lowest for the YNH participants and were higher for OMHL and OSNHL participants. While not plotted, the amount of masking was greatest for the YNH participants and progressively smaller for the OMHL and OSNHL participants. For the LFN masker, thresholds for the 25-ms masker-signal delay were significantly higher by 23 dB for the OSNHL than for the YNH participants (t=7.5, *p*<.001) but did not differ significantly for the OMHL and YNH participants (t=1.3, *p*=.700).

As hypothesized, threshold for the YNH participant group for the 25-ms masker-signal delay was significantly greater, by 8.5 dB, for the GN than for the LFN (t=5.1, *p*<.001), as also shown in Fig. 5. Consistent with the second hypothesis regarding expected effects of IHC saturation on GN disruption, GN disruption for the 25-ms masker-signal delay was significantly greater by 8.9 dB for the OMHL participants than for the YNH participants (t=3.5, *p*<.001). In contrast, GN disruption did not differ significantly between the YNH participants and the OSNHL participants (−1.5 dB, t=-0.6, *p*=.519). For the 75-ms masker-signal delay, GN disruption was statistically equivalent to that for the 25-ms masker-signal delay (−4.0 dB, t=- 1.7, *p*=.093) for the YNH participants. Note that this lack of a significant change in GN disruption may have been due to the larger GN disruption exhibited by two YNH participants for the 75-ms masker-signal delay (see the two outlier data points in Fig. 2).

GN disruption decreased significantly, by 8 dB for the YNH participants, from the 25- to 150-ms masker-signal delay (t=-3.4, *p*<.001). The decrease in GN disruption from the 25- to the 75-ms masker-signal delay was significantly greater for the OMHL than for the YNH participants by 10.2 dB (t=-2.9, *p*=.004). Similarly, the decrease in GN disruption from the 25- to 150-ms masker-signal delay was significantly greater for the OMHL than for the YNH participants, by 7.3 dB (t=-2.1, *p*=.040). The decreases in GN disruption from the 25-ms masker-signal delay to the 75- (t=0.3, *p*=.775) and 150-ms (t=0.9, *p*=.379) masker-signal delays were statistically equivalent for the YNH and OSNHL participants. Notice that variability in GN disruption, for the 25-ms masker-signal delay, was greatest for OMHL followed by YNH and OSNHL.

**Fig. 5.**
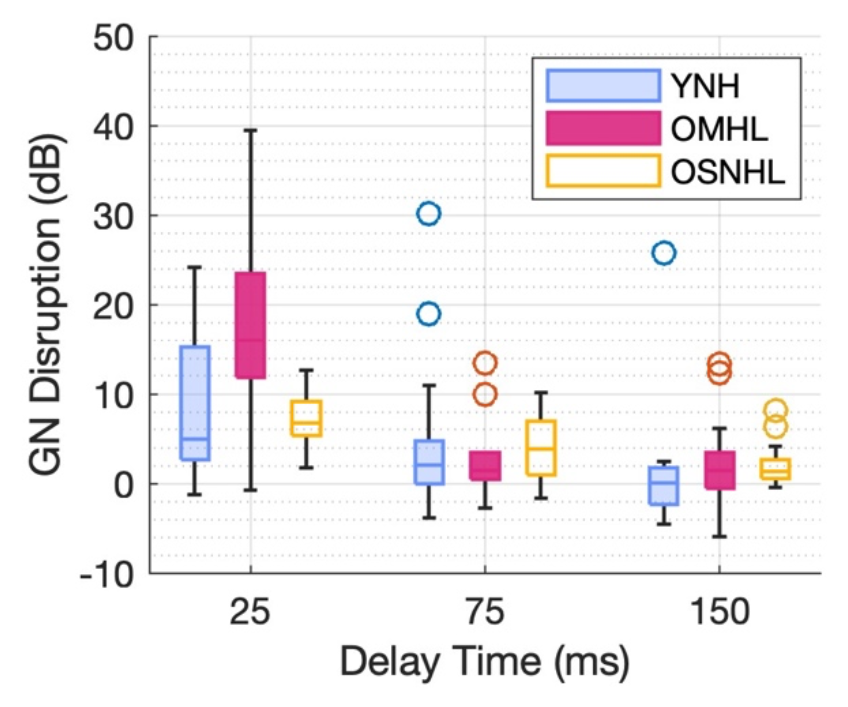
(color online) Plot of GN disruption. For the 25-ms masker-signal delay, GN disruption was larger for the OMHL participants than for the other two groups.

In summary, GN disruption for the 25-ms masker-signal delay was significantly greater for OMHL than for YNH and OSNHL participants. Recovery from GN disruption was greater (steeper) for OMHL than for YNH participants, with similar GN disruption for the three participant groups for the two longest masker-signal delays.

### C Computational Models

Figure 6 plots the simulated OHC gain as a function of time for each masker, masker-signal delay, and participant group. For all three groups and both masker types, OHC gain decreased following masker onset. For the computational model of YNH, OHC gain initially decreased due to an increase in the rate of the IC and LSR inputs to the MOC (see Fig. 2). Following this abrupt gain decrease, OHC gain gradually increased over the remaining duration of each masker. For the YNH group, the model predicts that these participants would likely be operating in IHC saturation in the presence of a masker presented at 80 dB SPL. Due to IHC saturation, the neural representation of input fluctuations of the GN masker would be reduced. The reduction of masker envelope fluctuations at the level of the IC results in a gradual OHC gain increase over the duration of each masker type. In contrast, for the computational model of OMHL, the model predicts that these participants would likely be operating below the saturation point of the IHC due to slight absolute threshold elevations, suggesting that envelope fluctuations from the GN masker would lead to strong neural fluctuations. This preservation of fluctuations at the level of the IC results in gradual OHC gain decreases, as opposed to increases (YNH), over the duration of the GN masker. Consequently, there were larger differences in OHC gain over time between the two masker types (GN vs. LFN) for the OMHL than for the YNH model. The OSNHL model predicts sensorineural hearing loss may lead to reduced OHC function and therefore reduced OHC gain increases or decreases, as well as reduced possibilities for IHC saturation. Consequently, smaller changes in predicted OHC gain were observed between the two masker types for this group. Notice too that the differences in OHC gain across groups also occurred during the signal for the 25-ms masker-signal delay. For all groups, the difference in gain at the time of the signal between the two masker conditions decreased for the longer masker-signal delays.

**Fig. 6.**
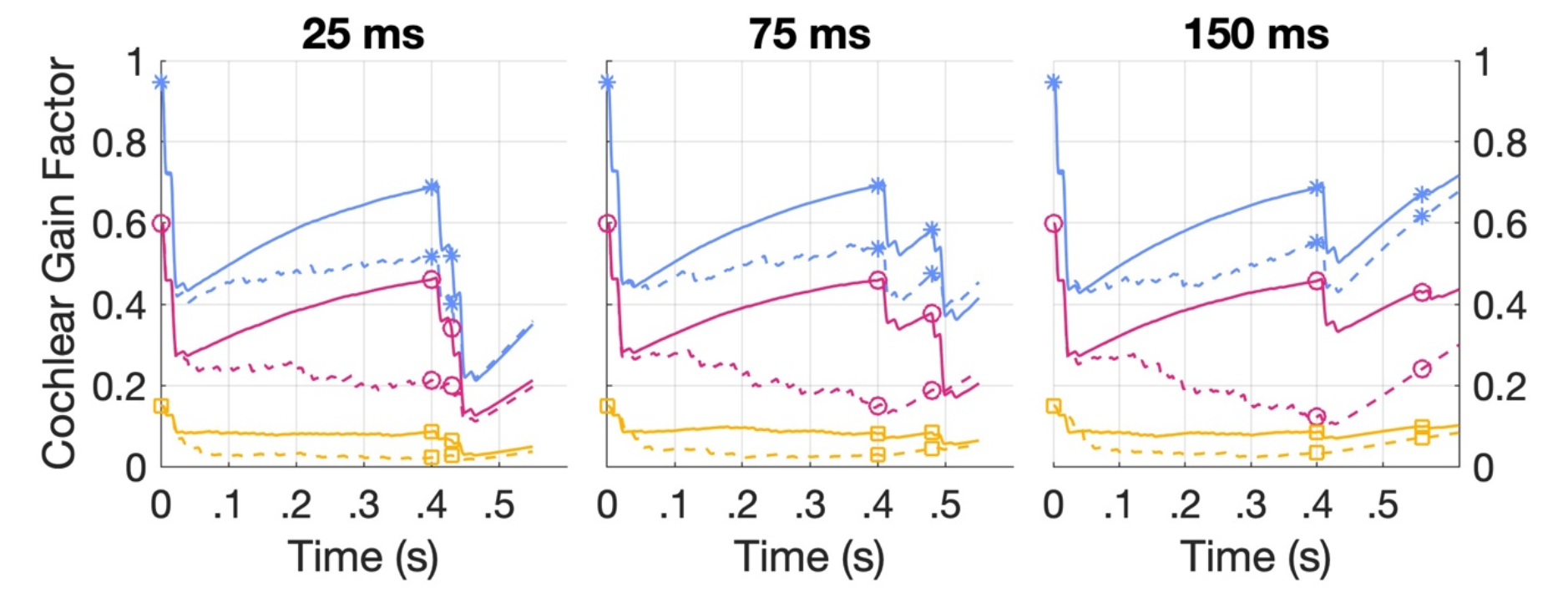
(color online) Plot of OHC gain for each masker-signal delay. Responses are shown for one hair cell with CF = 4 kHz. Differences in OHC gain between GN (dashed lines) and LFN (solid lines) were greater for OMHL (circles/magenta) than for YNH (asterisks/blue) and OSNHL (squares/yellow) models. Symbols denote, in temporal sequence, start of the masker, end of the masker, and temporal center of the 10-ms signal. The signal level was set 10 dB above the predicted model threshold for the GN conditions (see Fig. 7).

Figure 7 illustrates the masked thresholds estimated from the computational model, with and without the efferent system. As expected, masked thresholds with the efferent system were higher (poorer) than without the efferent system. The increase in thresholds for the GN relative to the LFN masker (i.e., GN disruption) was greater for the computational model when efferents were included. Note that the estimated thresholds were lower than the behavioral thresholds, with this difference largest for the YNH group. Figure 8 shows GN disruption estimated by the computational model, as well as measured behaviorally. Without the efferent system, GN disruption was minimal for YNH and OMHL. Starting with the 25-ms masker-signal delay condition, the estimates of GN disruption with the efferent system for YNH and OMHL groups increased by 4 and 18 dB, respectively, relative to the model without the efferent system. When the efferent system was included in the model, GN disruption was more similar in magnitude to the behaviorally measured GN disruption for all three participant groups. In addition, GN disruption estimated from the computational model with the efferent system was greatest for the OMHL group. GN disruption with the efferent system decreased for the two longer masker-signal delays and, with perhaps the exception of the 150-ms masker signal delay condition—better approximated GN disruption than the computational model without the efferent system.

**Fig. 7.**
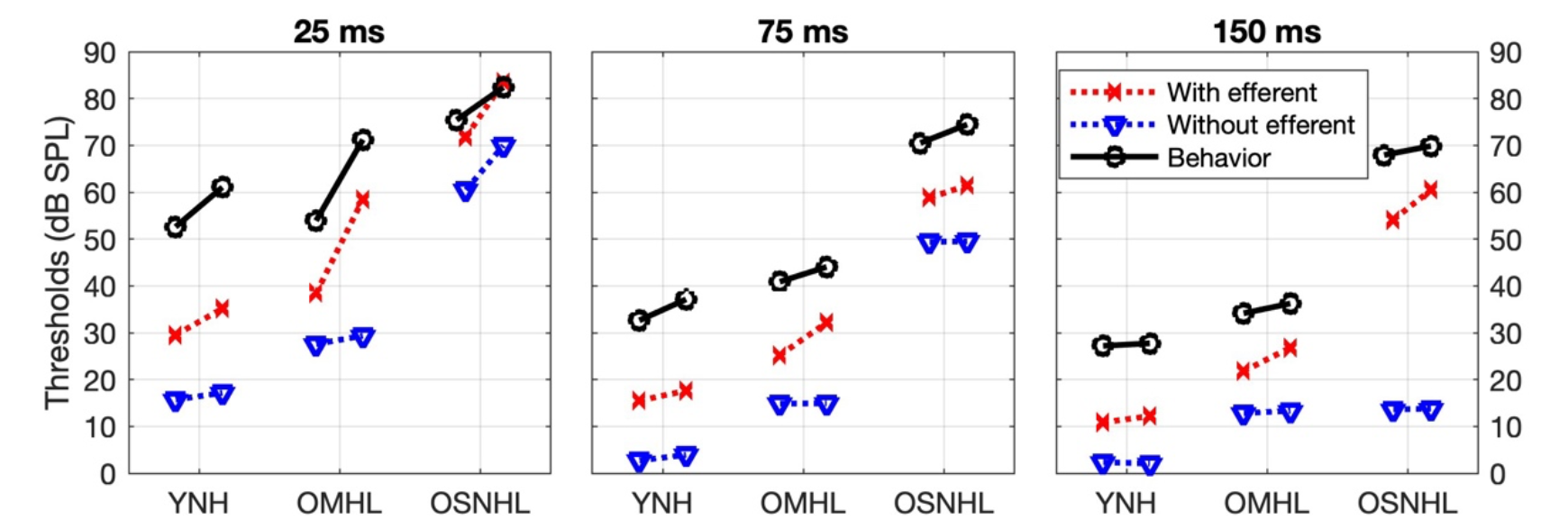
(color online) Plots of mean behavioral and model predicted thresholds for YNH, OMHL, and OSNHL with and without the efferent system. The masker-signal delay is indicated above each plot. For each participant group, LFN and GN thresholds are shown for the left and right symbols, respectively.

**Fig. 8.**
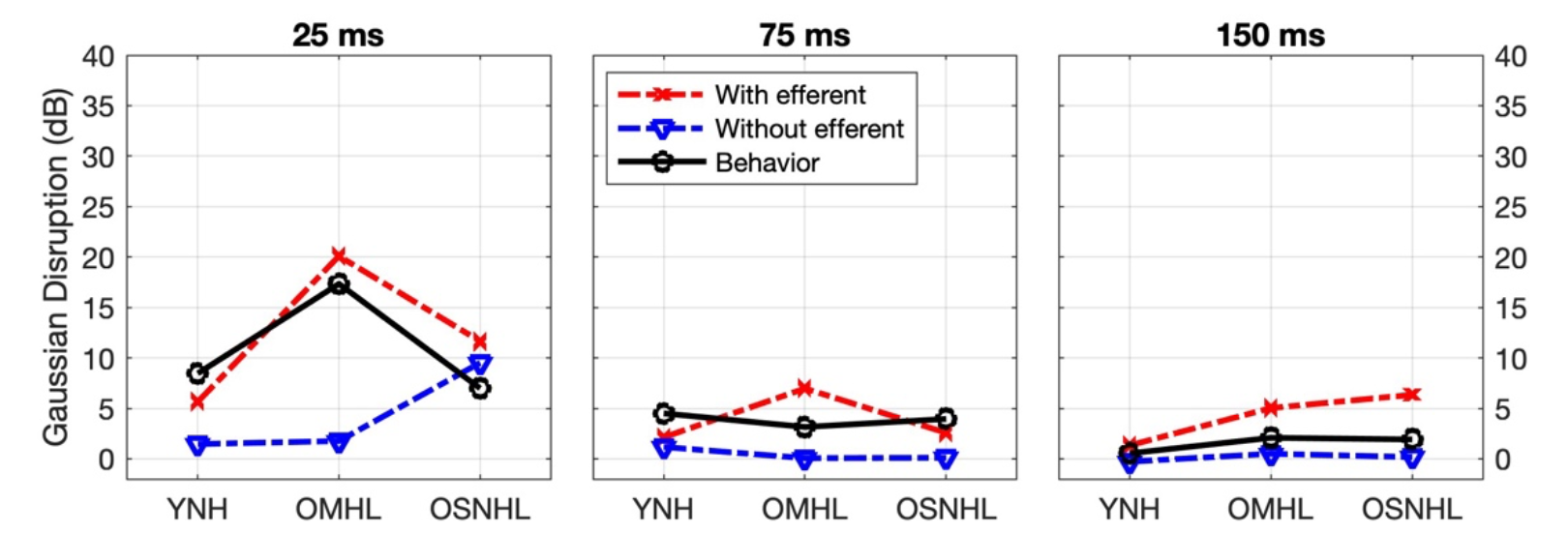
(color online) Plots of mean behavioral and model GN disruption for YNH, OMHL, and OSNHL with and without the efferent system. The masker-signal delay is indicated above each plot.

## IV DISCUSSION

### A Main Findings

Factors proposed to contribute to GN disruption include cochlear compression, listener uncertainty, inhibition within the auditory system, and the MOC efferent system. Because cochlear compression, inhibition, and MOC function vary with age and hearing status, this study was designed to: a) examine the contributions of each mechanism to GN disruption by comparing GN disruption for younger participants with NH to that for older participants with minimal hearing loss or SNHL; and b) examine the extent to which a computational model of the IC and MOC could account for the GN disruption observed in the behavioral results. The results suggest that the slightly elevated hearing thresholds of the OMHL participants may have made this group more susceptible than the other groups to the deleterious effects of masker envelope fluctuations. Specifically, greater GN disruption was observed for the 25-ms masker-signal delay for OMHL participants than for YNH and OSNHL participants and, based on the computational model, this greater GN disruption can be attributed to reduced IHC saturation. For all three groups, GN disruption was reduced for the two longer masker-signal delays (75, 150 ms), and at these longer delays, significant differences in GN disruption between the groups were not found.

### B Cochlear Compression

Prior investigators have hypothesized that linearization of the basilar membrane response associated with SNHL should result in greater effective amplitude fluctuations for GN, relative to a cochlea functioning normally (Moore *et al*., 1996; Svec *et al*., 2015). The greater effective magnitude of these fluctuations would likely cause increased difficulty for participants with SNHL, relative to the participants with NH, when attempting to detect a signal that follows a masker (Svec *et al*., 2016). If cochlear compression strongly contributed to GN disruption, then GN disruption should have been greater for the participants with SNHL than for the participants with NH. Instead, for the 25-ms masker-signal delay, GN disruption did not differ between the YNH and OSNHL participants and was *greatest* for the OMHL participants. While not statistically significant, these same trends were present in the GN disruption data of Svec *et al*. (2015). These results, at face value, do not support the argument that variations in cochlear compression across participants contributed to differences in GN disruption. However, it is possible that GN disruption for the OSNHL participants was a consequence of a quasi-linearized basilar membrane response with the absence of MOC efferent feedback. This notion is supported by the estimated GN disruption for the computational model without MOC efferent feedback. For this model, greater GN disruption (for the 25-ms masker-signal delay condition) was predicted for the OSNHL participants (9 dB) than for the YNH and OMHL participants (2 dB). This prediction of greater GN disruption for the OSNHL participants is consistent with a loss of OHC gain (and thereby compression) leading to larger fluctuations, and consequently, greater “modulation masking” by the GN masker.

### C Listener Uncertainty

While listener uncertainty, or confusion effects, have historically been observed at relatively brief masker-signal delays (< 20 ms; Moore 1981), Svec *et al*. (2016), showed significantly greater GN disruption for 10 OSNHL participants than for 9 YNH participants for a masker-signal delay of 75 ms, a finding that was not replicated here. Due to the lack of significant differences between groups, the results of the current study do not support the notion of larger GN disruption at longer masker-signal delays for older participants with minimal or greater hearing loss—as previously argued by Svec and colleagues. The computational model, on the other hand, makes a case for the involvement of the efferent system in what has instead been traditionally attributed to listener uncertainty. The effect of different temporal window durations on GN disruption was informally examined. Variations in the temporal window for which the window length changed from including to not including the masker significantly decreased GN disruption. Otherwise, variations in the temporal window had at most a minimal effect on GN disruption. Possibly, the participants with less GN disruption used shorter temporal windows that did not include the masker. Note that while listener uncertainty has been assessed using diotic presentation of a masker as a comparison to an entirely monaural stimulus presentation, binaural interactions at the level of the MOC and IC (Park, 1998, 2004) should perhaps give investigators pause. If, for example, cochlear gain is driven by both contralateral and ipsilateral projections from the MOC, then changes in masked threshold due to the introduction of a contralateral masker may not reflect solely a reduction in listener uncertainty but could instead reflect reduced cochlear gain from the MOC.

### D Inhibition and Envelope Coding at the Inferior Colliculus

The model simulations presented here suggest that forward suppression of the probe response is partially due to the effects of the MOC efferent system. This mechanism is different from offset-driven inhibition emerging in subcortical nuclei (Salimi *et al*., 2017). Simulations in the present paper demonstrate that the MOC efferent mechanism can explain different amounts of suppression for LNN and GN maskers. Whether offset inhibition could contribute to GN disruption remains to be examined. Note too that the computational model did not address changes in inhibition and envelope coding associated with age (Frisina and Rajan, 2005; Caspary *et al*., 2008; Parthasarathy *et al*., 2019), and thus, the authors cannot currently assess their contributions to GN disruption.

### E Role of MOC Efferent Feedback

Studies examining the physiological effect of MOC efferent feedback on firing rate within the IC provide supporting evidence that changes in rate within the IC may have contributed to GN disruption. The masker duration used here (400 ms) was sufficiently long to activate MOC efferent feedback (Cooper and Guinan, 2003) and the reduced GN disruption and forward masking for the longer masker-signal delays are consistent with the time course of OHC gain reduction following masker offset (Cooper and Guinan, 2003; Roverud and Strickland 2010; Guinan 2018). Note too that GN disruption, for some participants, was observed up to 150 ms following masker offset. Such an effect is consistent with the previously estimated 4 – 8 dB of cochlear gain reduction for a relatively long precursor-signal delay of 120 ms for some participants (Roverud and Strickland 2010). Together, these results provide evidence that reduction in OHC gain associated with the efferent system could have contributed to the GN disruption that was observed for the 25-ms masker-signal delay and, for some participants, up to the 150 ms delay.

Masked-threshold estimates from the computational model indicate there may be an influence of MOC efferent feedback on GN disruption. The computational model suggests that effects of masker type on firing rate within the IC without MOC efferent feedback are insufficient to explain GN disruption for individuals with normal hearing. For the 25-ms masker-signal delay, GN disruption estimated using the computation model without the MOC efferent system was only 2 dB for the 25-ms masker-signal delay, far less than the mean 8 and 17 dB measured for the YNH and OMHL participants, respectively. The addition of MOC efferent feedback to the computational model had the effect of reducing OHC gain within the model, which in turn, caused higher masked thresholds. More importantly, the amount of OHC gain reduction varied with masker type. There was a greater reduction in OHC gain for the GN than for the LFN masker. A greater reduction in gain likely occurred for the GN masker because the IC neurons that are excited by envelope fluctuations are assumed to excite MOC neurons, given that MOC neurons are also excited by envelope fluctuations (Gummer et al., 1988). As a result, for the YNH group and the 25-ms delay condition, OHC gain was reduced for the GN masker relative to the LFN masker (see Figs. 2 & 6), and this difference in OHC gain contributed to an additional 4 dB, for 6 dB total, of GN disruption relative to the computational model without MOC efferent feedback. Note too that the computational model with the efferent system predicted decreased GN disruption for the two longer masker-signal delays and these decreases in GN disruption were generally consistent with the decrease in GN disruption observed for the participants. In contrast, the computational model without efferents predicted smaller changes in GN disruption for the longer masker-signal delays that were inconsistent with the participant data.

The predictions of the computational model with the efferent system suggest that differences in the relative OHC gain over time between the three groups contributed to the differences observed in behavioral GN disruption between groups. Due to a loss of IHC sensitivity and because OHC gain started at a lower initial value (due to slight hearing threshold elevations), fluctuations within the IC were greater for the computational model for OMHL than for YNH. These more robust fluctuations caused OHC gain to *decrease* instead of increase over the duration of the masker. Consequently, a large GN disruption (for the 25-ms masker-signal delay) of 20 dB was predicted by the computational model, with 18 dB of the GN disruption attributable to the MOC efferent system. For OSNHL, the changes in OHC gain that occurred over time were markedly reduced relative to the other two groups, owing to their SNHL. Consequently, there was less difference in OHC gain between the two masker types, and therefore, only 2 dB of GN disruption was attributable to MOC efferent feedback.

Across participants, GN disruption ranged from −4 to 40 dB and this variability in thresholds was largest for the OMHL participants. The results of the computational model suggest that this variability in GN disruption can be attributed to variations in IHC saturation, which would be expected to increase with increasing minimal hearing loss and then decrease again for mild hearing loss. The smaller variance in GN disruption for OSNHL can be attributed to their reduced likelihood of IHC saturation in response to either masker type.

The current study contributes to prior work attempting to relate perception to MOC efferent feedback (e.g., Winslow and Sachs, 1988; Jennings *et al*., 2011; Wojtczak *et al*., 2019) by comparing masked thresholds obtained with a computational model of the midbrain with MOC efferent feedback to behavioral measures of GN disruption and provided evidence suggesting that the MOC efferent system contributes to GN disruption. Such an approach was also used by Jennings *et al*. (2011) to assess the role of MOC efferent feedback in overshoot, a phenomenon in which masked thresholds for a simultaneous masker are higher for a signal near the onset and offset of a masker than for a signal near the temporal center of a masker.

While the results of the computational model presented here indicate that modulation filtering, MOC efferent feedback, and IHC saturation provide plausible explanations for GN disruption, other aspects of auditory processing not assessed here could provide alternative explanations. The effect of several model parameters was not assessed, including the spontaneous rate of the fiber used for the decision variable and the best modulation frequency of the IC. In addition, the computational model did not capture all aspects of auditory processing, such as other cell types including sustained units (Krishna and Semple, 2000), the loss of auditory neurons associated with age (Makary *et al*., 2011), potential variations in the length of the temporal window across listeners or conditions or including multiple modulation-frequency channels. Possibly, some of these missing aspects of auditory processing in the computational model could account for 1) the better model thresholds as compared to behavioral thresholds and 2) individual differences in GN disruption.

Regarding combinations of different best modulation frequencies, the model thresholds for the NH listeners were elevated by inclusion of the efferent feedback, as expected. However, model thresholds were still about 25 dB lower than listeners’ thresholds. This gap might be addressed by a more comprehensive model for the MOC efferent feedback, for example, by including multiple modulation-frequency channels. The current model was based on a single band-enhanced IC model with a best modulation frequency of 64 Hz, based on the median of the distribution of best modulation frequencies in a representative mammal (Kim et al., 2020). Given that the mean modulation rate of the GN noise (98 Hz), incorporating modulation filters closer to the mean modulation rate of the GN noise could result in greater GN disruption. Future work may consider a model with additional modulation channels, perhaps spanning one or two octaves centered on 64 Hz, which would still fall within the range of modulation tuning observed in the IC and which would also pass significant modulation energy in response to the narrowband maskers used in this study. While one would expect such a model to elevate the predicted thresholds, especially in the model for YNH listeners, such an effect is not clear given that the modulation filters are broad (Q=1 and is consistent with data from rabbits as measured by Kim *et al*., 2020). The challenge for such future modeling efforts will be to design the combination of control signals across channels, as there is little physiological data to guide such model development.

## CONCLUSIONS

- It appears that IC sensitivity to masker fluctuations and the concurrent decrease in OHC gain contributes to GN disruption which has traditionally been attributed to mainly listener uncertainty.
- Participants with minimal hearing loss are more suspectable to GN disruption.
- A computational model of auditory processing in the IC with MOC efferent feedback indicated that increased saturation of IHCs, due to reduced efferent function, may have caused the greater GN disruption observed for the participants with minimal hearing loss.

Declarations of interest: None.

## AUTHORS CONTRIBUTIONS

MB and AS designed the research; MB performed the research; MB, BM, and AF analyzed the data and carried out the computational modeling; MB and AS wrote the paper. All authors reviewed the paper and provided feedback.

## ACKNOWLEDGEMENTS

Funding: This work was supported by the Nebraska Tobacco Settlement Biomedical Research Development Fund (MB) and the National Institutes of Health [010813 to LC, AF, & BM].

## Abbreviations

Auditory Nerve: (AN)
Auditory Nerve Fiber: (ANF)
characteristic frequency: (CF)
equivalent-rectangular bandwidth: (ERB_N_)
Gaussian noise: (GN)
inferior colliculus: (IC)
inner hair cell: (IHC)
low-fluctuation noise: (LFN)
low/high-spontaneous rate: (L/HSR)
Mean: (M)
Medial Olivocochlear: (MOC)
normal hearing: (NH)
outer hair cell: (OHC)
older participants with minimal hearing loss: (OMHL)
older participants with Sensorineural hearing loss: (OSNHL)
same-frequency inhibition-excitation: (SFIE)
sensorineural hearing loss: (SNHL)
younger participants with normal hearing: (YNH).

